# Continuous monitoring of glucose levels *in vivo* with a micro-organ based microfluidic biosensor

**DOI:** 10.1101/2024.04.03.587710

**Authors:** Emilie Puginier, Antoine Pirog, Florence Poulletier de Gannes, Julien Gaitan, Annabelle Hurtier, Marie Monchablon, Matthieu Raoux, Sylvie Renaud, Jochen Lang

## Abstract

Continuous monitoring of glucose levels has improved diabetes therapy. Current approaches rely on enzyme-linked electrochemical probes but do not allow a fully autonomous artificial pancreas. In contrast, monitoring the activity of a few electrogenic pancreatic islets in a biosensor may harness the computational power of the different endocrine cell types in the micro-organ, shaped for nutrient detection during evolution, and provide a more appropriate read-out. Extracellular electrophysiology captures slow potentials (SPs), which reflect coupled islet β-cell activity and is thus a method of choice for long-term monitoring of native islet activity *in vitro*.

We have now developed a microfluidic microelectrode chip containing a few islets and linked to interstitial fluids in live rats by subcutaneous microdialysis. The electrical activity in terms of slow potentials monitored by this biosensor reacts *ex vivo* proportionally to glucose levels off-line in serum or dialysed interstitial fluid. On-line monitoring *in vivo* reveals an excellent correlation between islet slow potential frequency, and to a lesser degree to slow potential amplitudes, to glucose concentrations with little variation between animals. The microorgan-based biosensor harness multiple parameters *in vivo* and provides a read-out closer to physiology. This demonstrates the usefulness of such biosensors for sensor-based therapy of diabetes.

## 1. Introduction

Pancreatic islets are at the centre of nutritional homeostasis and their demise cause the most common metabolic disease, that is type 1 or type 2 diabetes which both are characterized by increased blood glucose.^[1]^ Within the islet micro-organ, the insulin-containing β-cells function as actuator, which secrete insulin, the only glucose lowering hormone, as well as sensors, that measure the amount of nutrients available in the blood and thus turn food into a command for insulin release. An increase in ambient nutrient levels leads to an increase in β-cell metabolism and subsequently a change in ion channel activity and thus transmembrane ion fluxes.^[2]^ The ensuing depolarisation and calcium influx via voltage dependent ion channels induces highly regulated insulin release. The changes in ion channel activities are further regulated by hormones, eg the stress hormone adrenalin or enteric peptide hormones.^[3]^ Thus, transmembrane ion fluxes provide an integrative read-out according to nutritional status and to the state of the organism. Moreover, the final read-out in terms of insulin secretion is physiologically modulated by the other cell types present in the micro-organ, mainly α- and δ-cells.^[4]^

Type 1 diabetes is an autoimmune disease characterized by a large or complete loss of islet β-cells and requires hormone replacement therapy.^[5]^ As insulin, like most peptide hormones, exerts powerful action in the picomolar range, its concentration has to be carefully titrated to achieve therapeutic levels and avoid life-threatening hypoglycaemia. Previously, this was achieved via repetitive daily determinations of blood glucose levels by finger pricks. Recent technological advancements permit continuous blood glucose monitoring (CGM) via a subcutaneous electrochemical electrode.^[6]^ The use of CGMs has considerably reduced the discomfort for the patients and increased therapeutic precision as compared with previous discontinuous surveillance of glucose levels. This has led to the concept of an artificial pancreas where a subcutaneous sensor measures glucose in the interstitial fluid and commands a small insulin injecting pump as actuator via appropriate algorithms.^[7]^ Despite the progress achieved during the last 50 years, the system still does not work as a closed loop as it requires announcements of meals or physical activity and can provoke hypoglycaemia.^[6, 8]^

As pancreatic islets have been developed by half a billion years of evolution, they provide a fairly optimal sensor.^[9]^ In contrast to CGM they do not measure glucose but rather a “demand in insulin”, taking into account additional inputs such as amino-acids and incretin gut hormones. Furthermore, they contain internal inhibitory and stimulatory circuits given by the presence of the different cell types present in addition to β-cells.^[10]^ Moreover, in terms of activity they react stronger to a decrease in glucose than to an increase and this endogenous algorithm encoded by their electrical activity provides a clinical safety mechanism against hypoglycaemia.^[11a, 11b]^ Using these micro-organs as biosensor could thus offer considerable advantages in monitoring the nutritional state including glucose.

The capture of electrical cellular signals as signatures of activity offers a number of advantages as compared to other approaches. Gene transfer or chemical probes are not required for signal detection and the electronics dedicated to on-line signal analysis are well suited for miniaturization.^[12]^ We have previously studied and analysed in detail the electrical responses of human and rodent islets *in vitro* using extracellular electrophysiology with multielectrode arrays, a non-invasive method that allows recording over long time periods.^[11b, 13]^ The recorded slow potentials (SP) are generated by the multicellular islets and their amplitudes provide a direct read-out for coupling between β-cells, a physiological hallmark of islet β-cell function.^[14]^ These electrical signatures of islet activity, as recorded *in vitro* by MEAs, can be introduced in a simulator of human metabolism in T1D patients, called UVA/Padova.^[15]^ This computer model simulates the glucose-insulin dynamics in T1D patients, and is approved by the US Food and Drug Administration (FDA) as an alternative for pre-clinical testing of insulin therapies, including closed-loop algorithms. Within this *in silico* model, an islet biosensor-based artificial pancreas was as efficient as standard CGMs and even outperformed them under challenging conditions.^[16]^

We therefore asked whether monitoring the electrical activity of a few islets may provide a sensor for continuous glucose measurements in live animals. To this end we developed stepwise microfluidic micro-electrode arrays containing a few islets and interfaced with the interstitial fluid via microdialysis. Our data and their analysis indicate faithful monitoring of glucose levels in rats during glucose tolerance tests and subsequent to insulin injections.

## 2. Results

### 2.1. Experiment design

A biosensor using heterologous live cells, such as pancreatic islets, has to be conceived as an extracorporeal device and microdialysis can provide continuous access to bodily fluids such as interstitial liquids. As the amount of interstitial fluid that can continuously be retrieved is limited, miniaturisation is required. To test an islet-based biosensor for continuous nutrient monitoring, interfacing of the animal with the sensor was developed as given in **Figure 1**. Interstitial fluid is obtained via a microdialysis pump and a subcutaneous microdialysis catheter which is linked to a chip consisting of the sensor, a microelectrode array with islets attached to its electrodes and the microfluidic system to pass the dialysate to these islets (Figure 1A). To assess the biosensors’ characteristics, recordings have to be compared with blood glucose, which can be measured after small incisions at the rat’s tail vein and dialysate was also sampled after passage through the chip to determine its glucose concentration. This configuration has to deal with a number of delays between blood glucose and the sensor. First, diffusion of glucose in the interstitial space requires some 10 minutes in man and rat.^[17]^ Second, in this laboratory set-up a certain length of tubing is required to link the components introducing additional delays between the point of dialysis and the microfluidic MEA (µMEA) as well from the µMEA to the point of glucose measuring in the dialysate at the outlet (Figure 1B). We have tested these delays using phenol red as dye in the fluids and the resulting delay time (Figure 1B, see also Methods) was used throughout when comparing different parameters on a time scale. Figure 1 C to E shows the actual set-up on the bench with a video-microscope to observe potential formation of bubbles in the microfluidic channels (Figure 1C), the microfluidic MEA (Figure 1D) a rat with an implanted interscapular catheter for microdialysis (Figure 1D).

**Figure 1:**
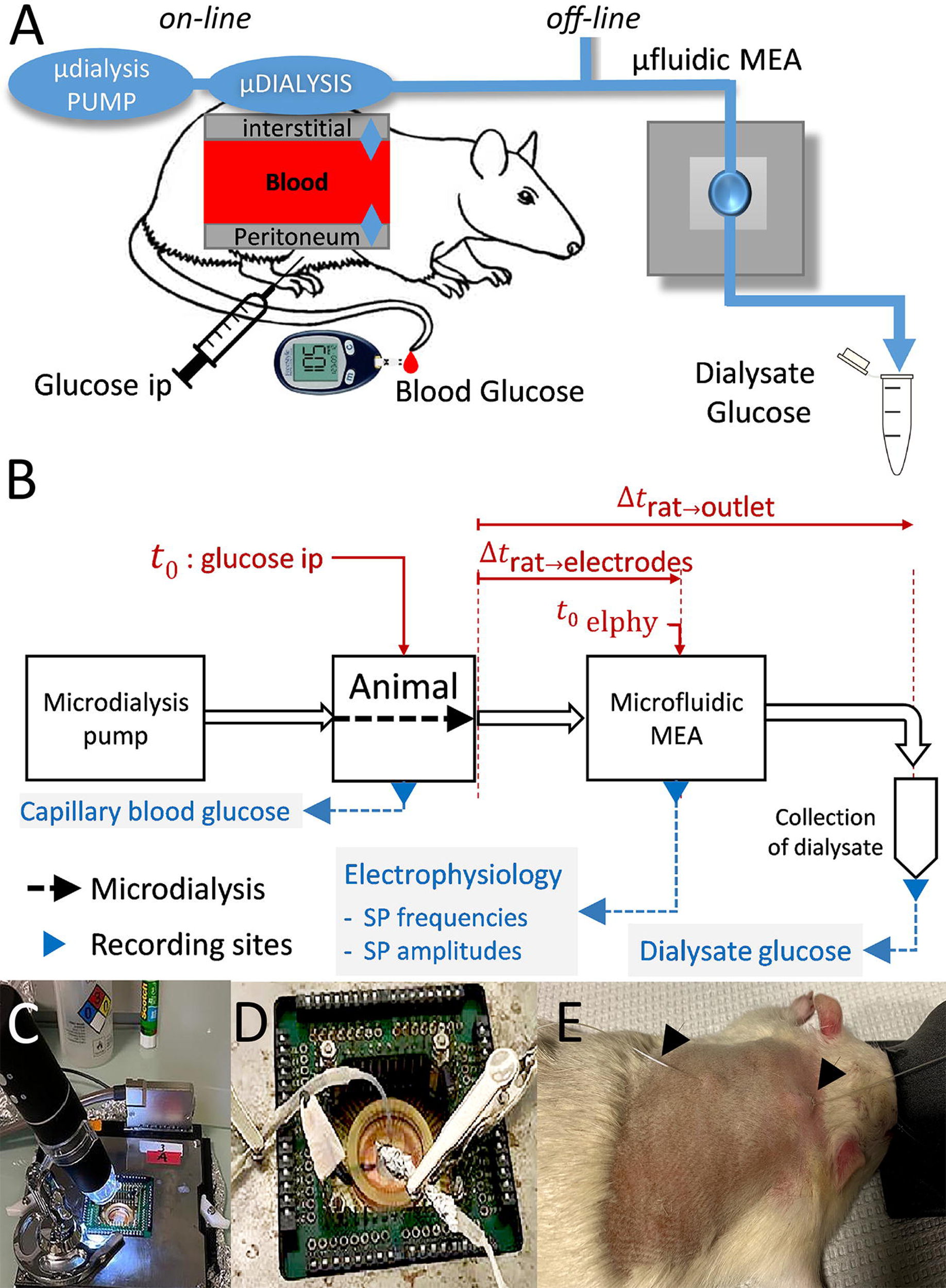
Experiment design. **A**: Anaesthetized rats were subjected to an intraperitoneal glucose tolerance test and blood glucose was determined. In off-line experiments, serum samples were added directly to the microfluidic MEA; in on-line experiments, interstitial fluids were dialyzed at 1 µl/min and fed to the microfluidic MEA. Glucose concentrations in the dialysate were determined off-line from an outflow channel of the microfluidic MEA. **B**: Full set-up and work flow for on-line experiments. Time points in the experiment and delays introduced by tubings between microdialysis and the different recording sites are given. Δt_rat-electrodes_ was 750 s, Δt_rat-outlet_ (collection of dialysate) was 2460 s as determined by video films using a dye (see Methods). **C**, MEA setup with microscope video camera to inspect flow. **D**, enlarged view of the MEA itself and microfluidic inlet/outlet. **E**: anesthetized rat with dorsal subcutaneous catheter inserted, arrows at inlet and outlet of subcutaneous dialysis.

### 2.2. *Ex vivo* monitoring of serum glucose and characterization of subcutaneous microdialysis

The final device was interfaced gradually. Previous work had used defined electrophysiological buffers and revealed the presence of two different electrical islet cell signals that can be recorded by extracellular electrophysiology; single cell action potentials that are difficult to capture due to their minute amplitude, and robust slow potentials (SP) that are generated by cell to cell coupling. Their amplitude depends on the degree of coupling between β-cells, which is hallmark of physiological islet function.^[11b, 18]^

As islets on MEA had never been challenged with sera, we first tested the response to human or rat serum containing different glucose concentrations in static incubation in MEAs with home-made PDMS microwells to allow assaying of a few microliters of analyte. As shown in **Figure 2** (A, B, E), islets exhibit strong responses in terms of SP frequency and amplitude in the presence of culture medium containing 11 mM glucose and amino-acids. Replacing medium by human serum (6 mM glucose) induces a rapid decrease in electrical responses. Subsequently islets were exposed sequentially to human serum containing 9, 12 or 15 mM glucose. Statistical evaluation of the corresponding areas under the curve (AUCs) revealed significant differences in activity between all the glucose ramps from 6 to 15 mM (Fig 2 C, F). A high degree of correlation between glucose concentrations in human serum and recorded responses was evident (Figure 2 D, G) and suggested a good discrimination power of the biosensor. In a next step, we performed an intraperitoneal glucose tolerance test in a rat with a basal glucose level of 9.4 mM. Injection of 2 mg/kg of glucose led to a transient increase in blood glucose (Figure 2H) to 22.9 mM followed by a slow decrease to 11.9 and 8.6 mM. At each determination of blood glucose, sera were prepared and added ex tempore to PDMS microwells fixed on MEAs (Figure 2 I, K). The change from 9.4 to 22.9 mM glucose induced a strong response in the biosensor in terms of SP frequency and amplitude. Responses were clearly distinct among the conditions and were all significantly different for SP frequencies and also for SP amplitudes except for G12 vs G9 in the latter case (Figure 2, J and L).

**Figure 2:**
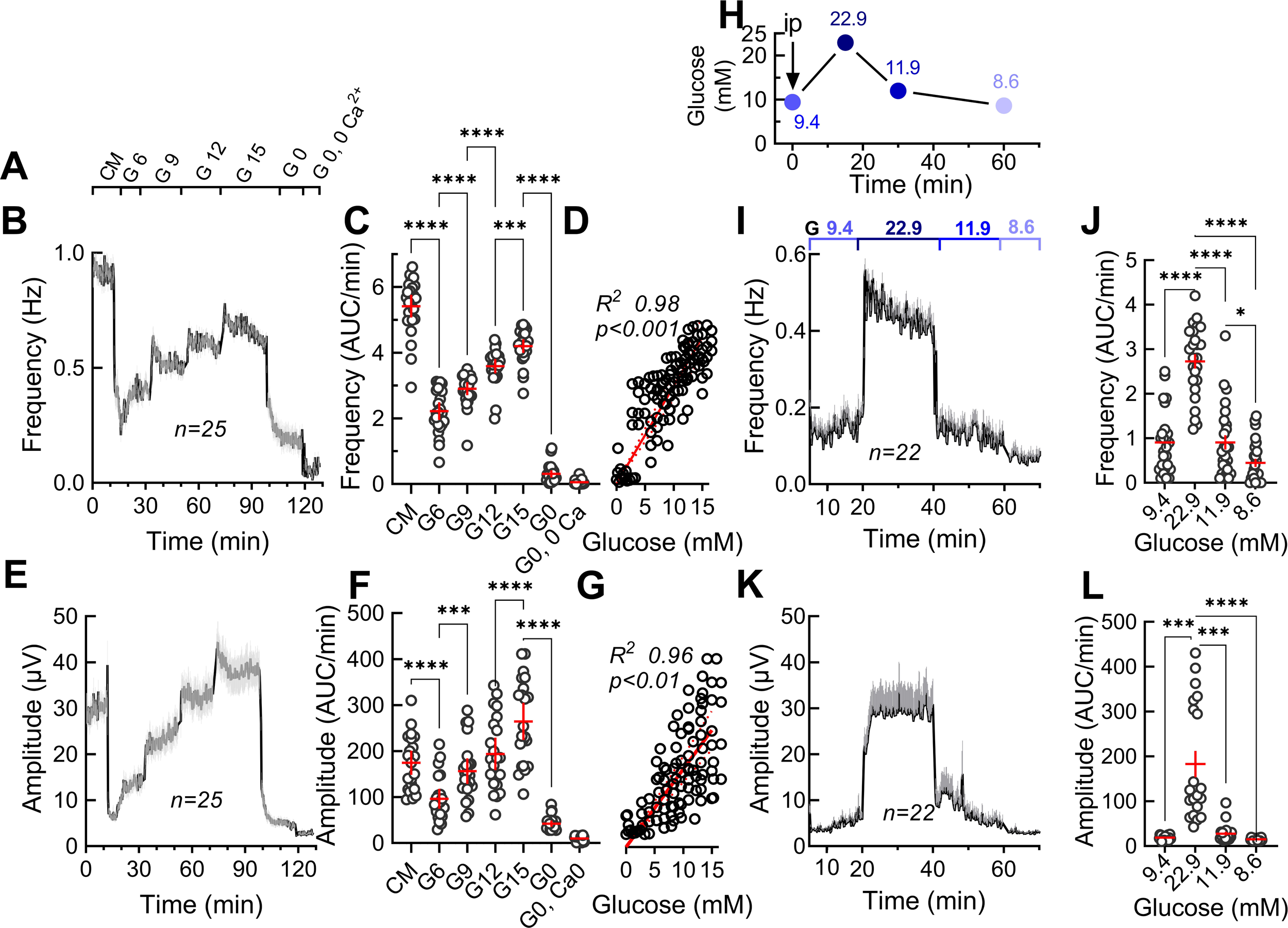
*Ex vivo* monitoring of human or rat serum glucose. **A:** Solutions used; CM, culture medium containing amino acids and 11 mM glucose; G6, human serum containing 6 mM glucose; G9 to G15, same human serum adjusted to 9, 12 or 15 mM glucose; G 0, buffer without glucose; G0, 0 Ca^2+^, buffer without glucose and calcium. **B**: Frequencies of slow potentials recorded by microwell MEAs under the different conditions given in A. **C**: Areas under the curve from B determined during the first 10 min of each stimulus and expressed as AUC/minute. **D**: Pearson correlation analysis of C (SP frequency vs. glucose concentrations). **E**: Amplitudes of slow potentials recorded by microwell MEAs under the different conditions given in A. **F**: Areas under the curve from E determined during the first 10 min of each stimulus and expressed as AUC/minute. **G**: Pearson correlation analysis of C (SP amplitude vs. glucose concentrations). **H:** Intraperitoneal glucose tolerance test in an anaesthetized rat. Blood samples were collected at indicated time points, corresponding sera prepared and glucose concentrations determined. **I:** Effect of rat sera in on slow potential frequencies in microwell MEAs. Glucose concentrations in the sera are given on top. **J**: statistical analysis of AUCs determined as in I. **K:** Effect of rat sera on slow potential amplitudes during same recordings as in G. **L**: statistical analysis of AUCs determined as in C. Means and SEM are indicated in black and grey in B, I, E and K, and in red in the other panels. *, 2p<0.05; ***, 2p<0.001; ****, 2p<0.0001 as compared to the presence of different glucose concentration. ANOVA and paired Tukey post-hoc tests, n as given in the corresponding panels.

Having thus validated that the islet-based biosensor can be used with serum and provides discrimination between glucose levels differing by a few millimoles/liter, we tested rat microdialysis dialysate off-line with the MEA biosensor (**Figure 3**). Blood glucose recovery by dialysis at a fixed glucose level depended on flow rates (Figure 3A). The rate of 1 µl/min provided sufficient glucose recovery from a test sample (Figure 3A) and a reasonable flow. This setting was chosen for all subsequent experiments and provided a constant recovery at about 78% at low and high glucose levels during a glucose tolerance test in rat (Figure 3B). Next we tested the two dialysates obtained in rat before (t=0) and during the glucose tolerance test (t=30) in the biosensor. We observed a strong increase in activity when changing from 5.8 (t=0) to 15 mM (t=30) dialysate glucose in terms of frequencies (Figure 3C, D) and of amplitudes (Figure 3E, F). Notably a first and second phase was visible especially in terms of frequencies and amplitudes (Figure 3C, E), which is due to differences in coupling during islet β-cell activation.^[13c]^

**Figure 3.**
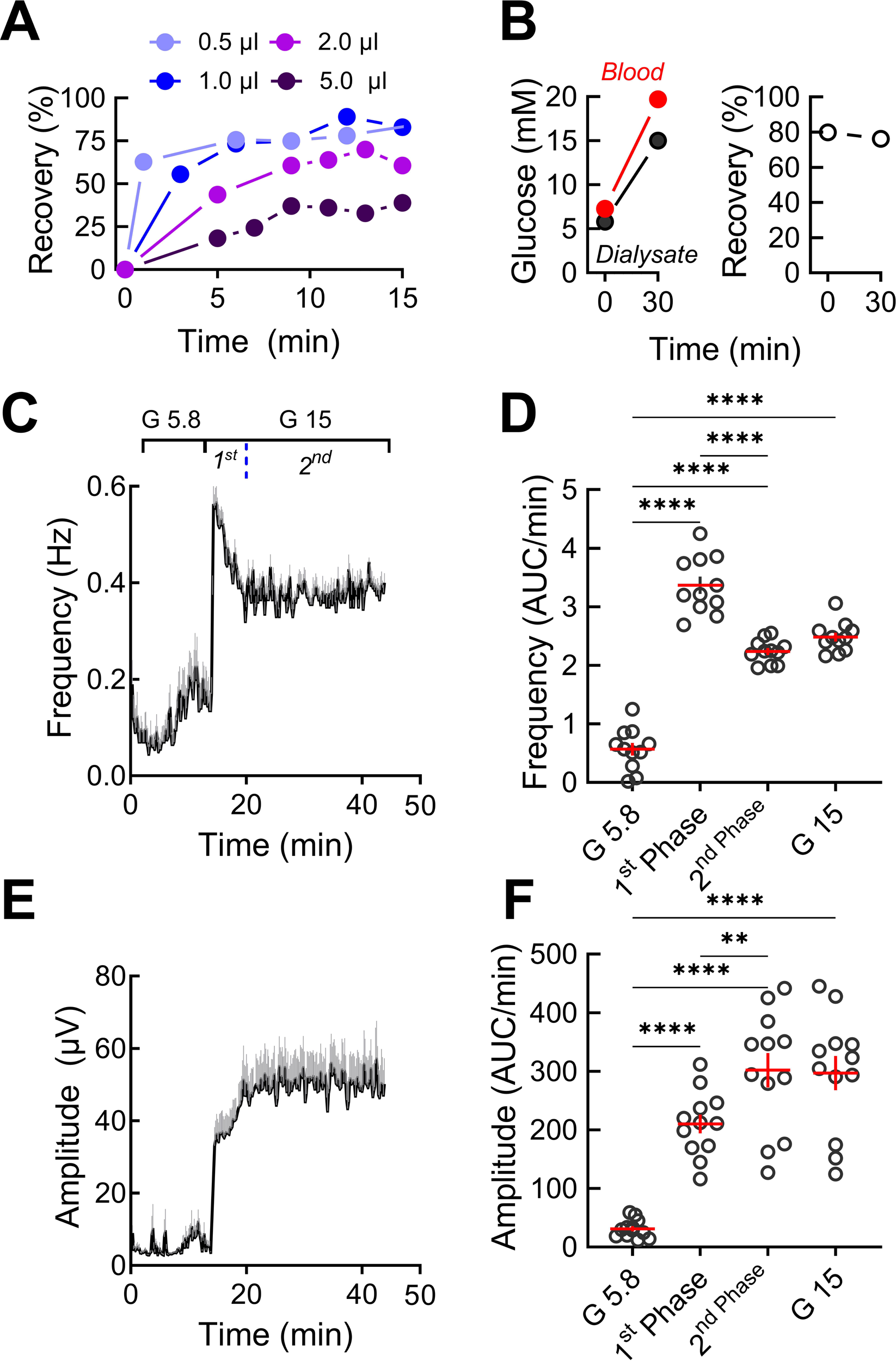
Characterization of subcutaneous microdialysis and dialysate effect on islet electrical activity. **A:** Recovery rates for glucose in saline buffer at different flow rates per minute (0.3-2 µl/min). **B:** Glucose concentrations in rat capillary blood and in dialysate (flow rate 1 µl/min) after intraperitoneal injection of glucose (ip) and recovery rates **C:** Effect of corresponding dialysates (at 5.8 and 15 mM glucose) on islet slow potential frequencies. Blue dotted line indicates 1^st^ vs 2^nd^ phase. **D:** Statistics of areas under the curve (G 5.8, 0-14 min; 1^st^ phase 14-18 min; 2^nd^ phase 20-43 min; G15 14-43 min; expressed as AUC/min). **E:** Effect of corresponding dialysates (at 5.8 or 15 mM glucose) on islet slow potential amplitudes. **D:** Statistics of areas under the curve (details see D). **, 2p <0.01; ****, 2p<0.0001 (Tukey post-hoc test), n=12.

### 2.3. Characterization of microfluidic micro-electrode arrays

We initially explored configurations allowing single islet trapping on metal electrodes to accommodate the small flow rates of microdialysis. Although such a set-up worked satisfactorily *in vitro*, this was not the case when interfacing this with microdialysis and a live animal despite all diligence and bubble traps. Moreover, islets are known to be heterogeneous in size and relative occurrence of the different endocrine cell types.^[19]^ Therefore, recording from a number of islets and not just single islets provides a more physiological readout. We therefore opted for a simpler and more robust approach (**Figure 4**) consisting of a PDMS block with a single channel, inlet and outlet (Figure 4A) that was aligned on half of the 60 electrodes of a commercial MEAs in a cleanroom under a binocular microscope as shown in Figure 4B. The microfluidic channel is charged with islets that cover a reasonable number of electrodes and stay in good shape (Figure 4C). The simultaneous recording from multiple islets also ensures a reasonable statistical sampling. Simulations of flow dynamics revealed shear stress mainly at the bottom of the channel where islets are adhering but the maximal value of 250 µPa predicted at a flow of 1 µl/min remains still below values that have been reported as critical.^[20]^

**Figure 4:**
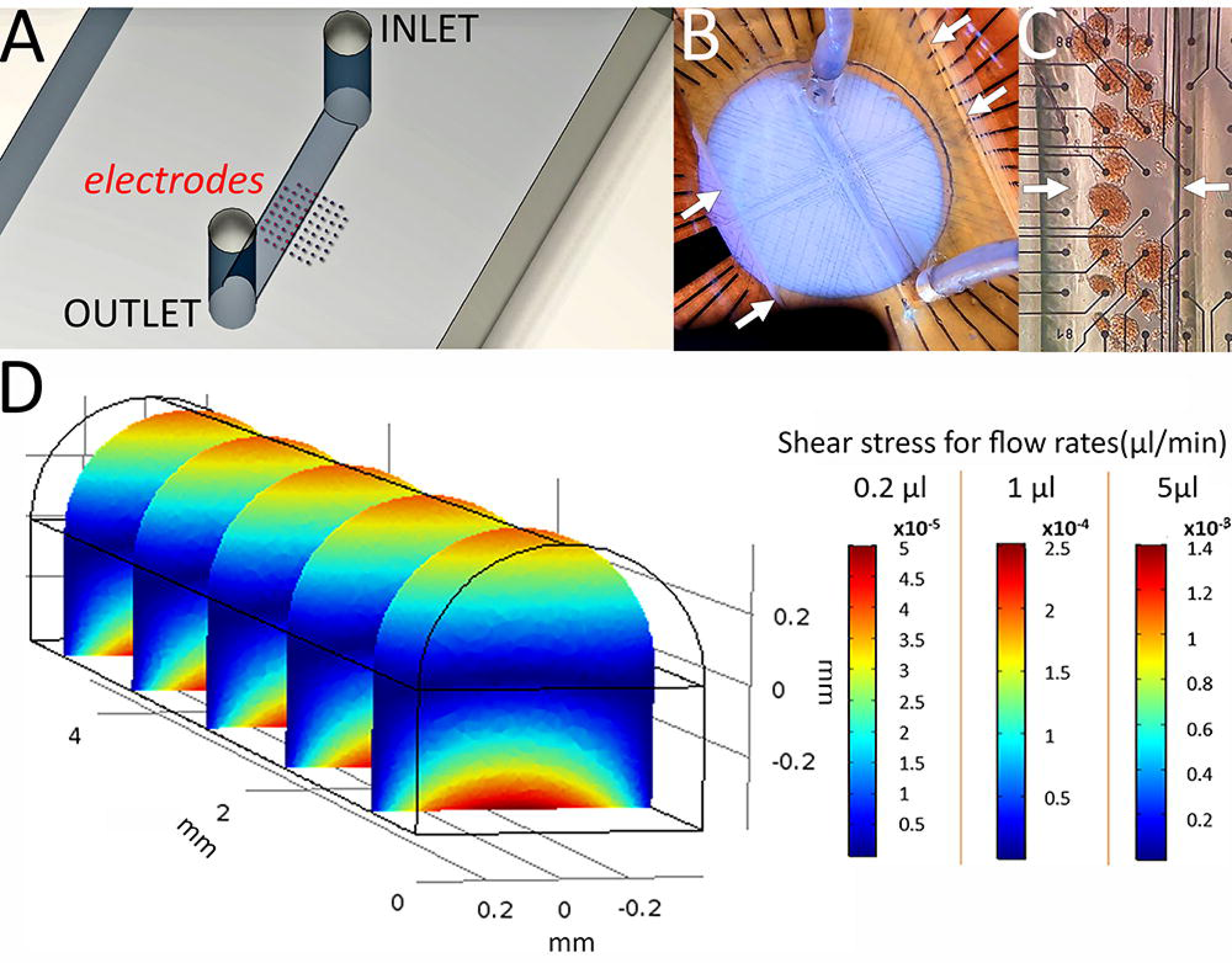
Characterization of microfluidic micro-electrode arrays. **A:** Layout of microfluidic MEA. **B:** View of a microfluidic PDMS device on a MEA, inlet and outlet are visible. Arrows, border of the PDMS device. Channel length 10 mm, diameter 0.8 mm. **C:** Zoom on the electrodes aligned in the microfluidic channel of the chip Arrows, lateral borders of the channel. **D.** Multiphysics simulations provide a rainbow view of the shear stress as z axis repartition over the channel at a flow-rate of 0.2, 1 or 5 µl/min. Scales on the right show the color codes of the values reached for the different flow rates simulated (color codes were adapted scaled here for different flow rates to apply all for the simulation shown on the left).

### 2.4. *In vivo* monitoring of glucose concentrations

We coupled this optimized µMEA to microdialysis to compare electrical activity profiles of the biosensor with blood and dialysate glucose values as an indicator for its potential usefulness in continuous nutrient monitoring. As an independent control of the condition of the animal we controlled respiration rates as breathing may influence the performance of the interscapular microdialysis catheter and could also introduce mechanical artefacts on the biosensor device. As given in **Figure 5A**, respiration rate was stable and in a normal range of 1 Hz throughout the entire procedure. After a first glucose injection, we observed a rapid increase in blood glucose which reached its maximum after 40 to 60 min, whereas dialysate glucose was retarded by around 10 min. Both measures were discontinuous as the low microdialysis flow rate requires a minimal collection volume and time for subsequent glucose determination. The increase in glucose was mirrored by an increase in electrical activity in terms of slow potential frequency and amplitude and a peak was attained at 60 to 70 min followed by a decrease in line with glucose concentrations. Note that, as expected, the slow physiological increase in glucose here did not induce a biphasic electrical response in contrast to the clear presence of biphasic pattern after stepwise increases in glucose (Figure 2B and 3C). To test the reactivity of the system, insulin was injected and as expected blood as well as dialysate glucose levels decreased correspondingly. Subsequently to both injections of insulin a marked decrease in electrical activity was apparent (Figure 5C, D).

**Figure 5:**
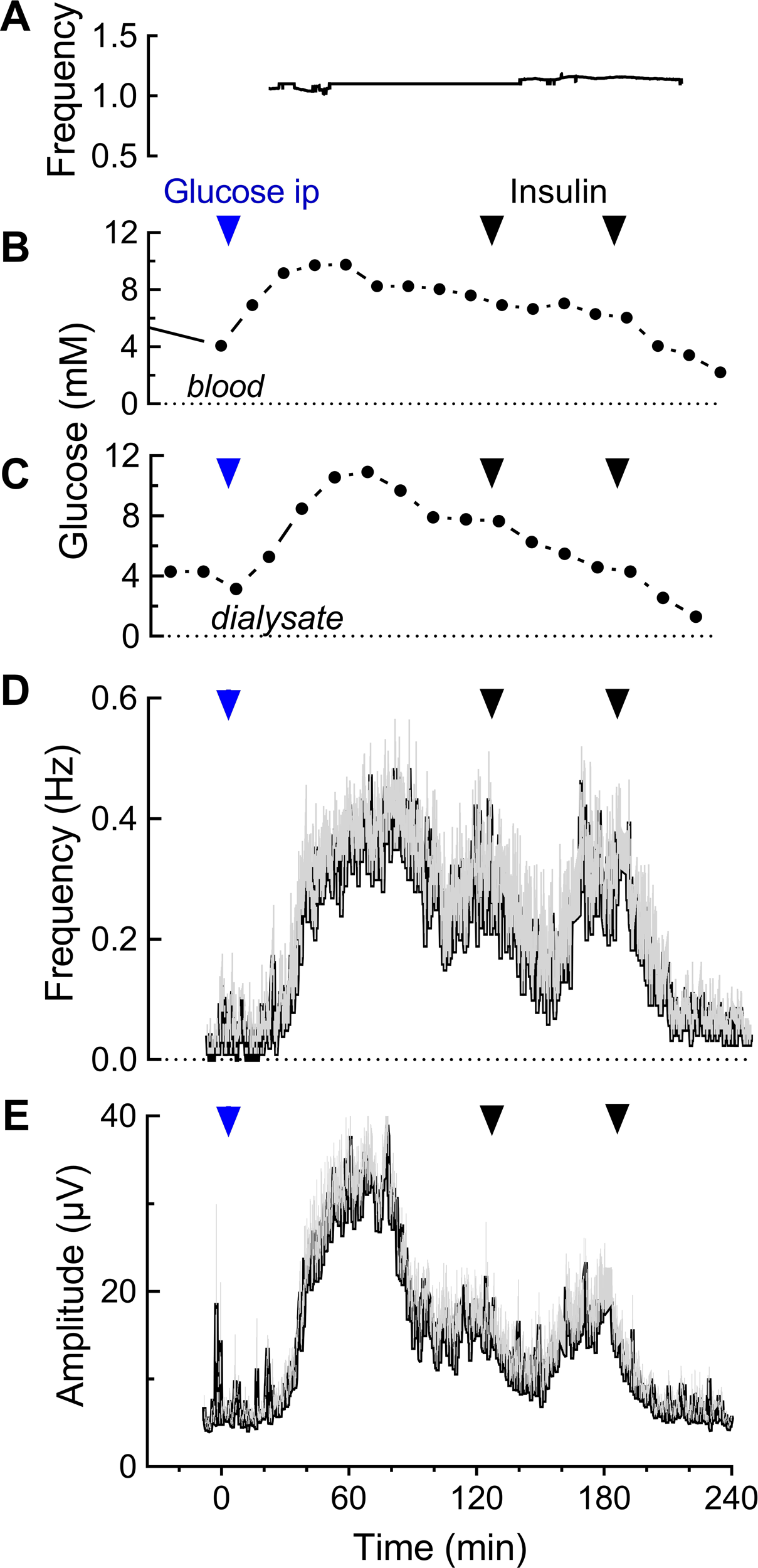
*In vivo* monitoring of blood or interstitial glucose concentrations and dialysate-evoked islet cell activity in the microfluidic MEA device. The time scales given here take into account the determined delays between microdialysis and microfluidic MEA (electrical recordings) as well as MEA outlets where glucose was determined in the dialysate (see Fig. 1). **A:** Respiration frequency (per minute) of anaesthetized rat. **B:** Blood glucose levels, arrows indicate intraperitoneal glucose (blue) injection or subcutaneous insulin injections (black). **C:** Dialysate glucose levels as measured after its passage through the microfluidic MEA. **D:** Islet electrical activity in terms of slow potential frequencies. Arrows indicate time of glucose or insulin injection, n=6. **E:** Islet electrical activity in terms of slow potential amplitudes, n=6.

In contrast to *in vitro* systems, where precise concentrations can be imposed, the absolute changes and kinetics in glycemia vary among animals. Moreover, the electrical activity of the biosensor islets is monitored at a microsecond scale, whereas blood or interstitial glucose is measured at far greater intervals. To obtain insight about the correlation between blood or interstitial glucose and biosensor responses we calculated the AUCs of electrical responses over the same time span as the fluid collection span for four (dialysate glucose) or five (blood glucose) independent experiments to compare equal time spans. Linear regression analysis for each independent experiment provided a set of graphs that were highly parallel in the case of SP frequencies whereas variable slopes were observed for amplitudes (**Figure 6A, B**). Analyses for dialysate glucose are given in Suppl. Figure 1 and a comprehensive view of Spearman’s ρ values is provided in **Figure 6C**. Each of these correlations for a given experiment was highly significant (Supplementary Table 1). The biosensor response in terms of frequencies versus amplitudes was highly correlated as were electrical activities (frequencies and amplitudes) versus blood glucose levels with means around 0.9. The correlation between electrical responses and dialysate were more scattered which may be due to its measurement after passage through the microfluidic MEA and potential diffusion phenomena especially at low flow rates. However, they were still highly significative for association with Spearman’s ρ values mean values between 0.8 and 0.9 (**Figure 6C** and Supplementary Table 1). The scatter plot in **Figure 6D** shows the coefficient of determination of linear regressions and the identified slopes, which measure the linearly dependent nature of the studied metrics regardless of basal values. The spread of values on either axis helps visualize the homogeneity of the results across experiments. As such, the clustering of our experimental values in the scatter plot reflects the repeatability of the fold increases in the studied metric relative to basal conditions. The data indicate a more homogeneous distribution of effects on frequency as compared to amplitudes.

**Figure 6:**
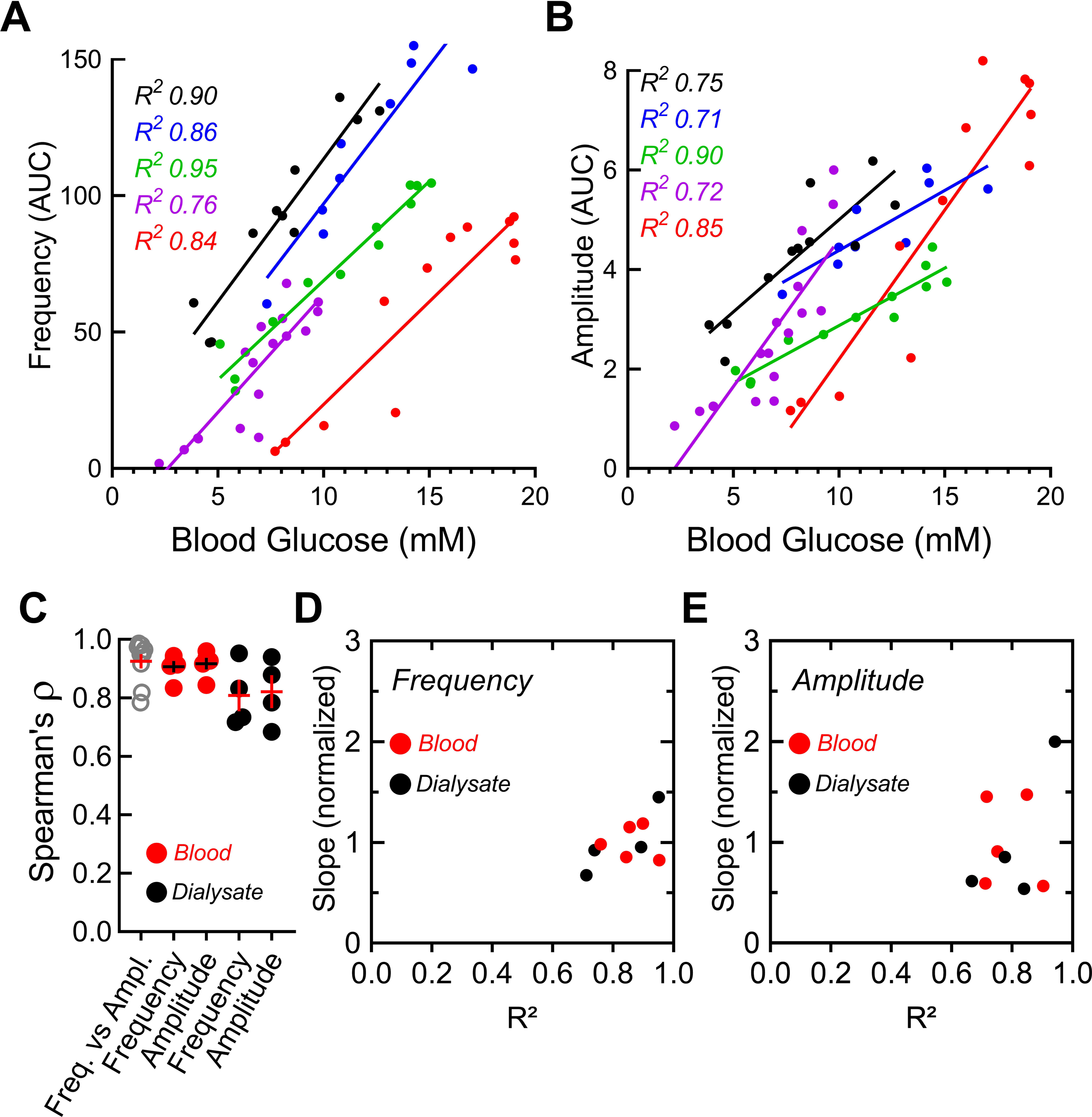
Correlation between electrical signal and blood and dialysate glucose levels. **A:** Linear regressions between glucose levels and slow potential frequencies. Each color represents one experiment (animal) and corresponding coefficients of determination R^2^ are indicated. Data points represents AUCs for a given blood glucose value and regression curves are given. **B:** Linear regressions between blood glucose levels and slow potential amplitudes. Color codes as in A, corresponding R^2^ values are indicated. **C:** Spearman correlation for slow potential frequencies vs amplitudes over the blood glucose range during experiments (grey) as well as for frequency or amplitudes versus glucose values either in blood (red) or in dialysate (black). Each data point represents one experiment (animal), mean and SEMs are given. **D** and **E:** Distribution of slopes vs R^2^ values for slow potential frequencies or amplitudes. R^2^ and slope values are from regression analyses between electrical signals and blood or dialysate glucose levels, each point represents one experiment (4-5 animals, each 2 – 13 electrodes).

### 3.0 Discussion

Continuous monitoring of nutrient levels remains a major challenge in diabetes therapy and despite a remarkable progress over the last 50 years, an autonomous closed loop system is still not available.^[6–7]^ Our work here demonstrates the feasibility of a label free microorgan-based biosensor and its precise recording of vital parameters.

Regression analysis over a number of *in vivo* experiments indicates a remarkable homogeneity in terms of sensor responses to different glucose levels despite the variability in biological systems. In each experiment different animals as well as different preparation of islets for the biosensor were used and intrastrain variation in metabolic responses is well known.^[21]^ As frequency and amplitudes of slow potentials were analysed, the question arises whether one or the other quality is closer related to glucose levels. Previous *in vitro* experiments suggested that slow potential frequencies reflect more closely insulin secretion and is further refined when taking also amplitudes into account.^[13c]^ However, *in vitro* experiments use strong square shaped stimulation by sudden increase in glucose and provoke a biphasic response. Such a biphasic response was apparent here only upon sudden increase by externally applied steps of 9 or more mM glucose but not during slow increases when coupled to microdialysis. The existence of biphasic activity and secretion *in vivo* in man is debated and may be absent during absorption of a meal.^[22]^ Our data suggest that a frequency-based analysis may be more robust during *in vivo* applications. Interestingly, the biosensor reacted strongly to insulin injection despite minor concomitant decreases in blood and dialysate glucose. It is known that glucose dependency of insulin secretion displays a hysteresis when comparing increases versus decreases in glucose levels in man *in vivo*.^[11a]^ This serves as a kind of safety break to avoid hypoglycaemia and is also found in isolated islets *in vitro*.^[11b]^ Although we have not investigated the existence of a hysteresis here, one may speculate that the considerable decrease upon insulin injection observed here may be due to such an islet mechanism.

We have also projected a possible packaging of an islet-based biosensor (Supplementary Figure 2) including microdialysis and an insulin reservoir. Although the qualities of a micro-organ biosensor are evident, there are also limitations and obstacles. In contrast to enzyme-based electrochemical sensors, microdialysis is necessary with concomitant space requirements and device duration. Note, however, that an electrochemical sensor linked to microdialysis had been commercialized previously and successfully used in humans.^[23]^ As islet encapsulation techniques are constantly evolving, direct implantation of such a device may eventually be possible in the future.^[24]^ A clear limitation is given by the type of islets to be used in the sensor. Reaggregated human donor islets exhibit excellent function but may raise ethical issues by diverting islets from use in transplantation.^[25]^ Alternatively, stem cell-derived β-cell like cells have attained considerable functional maturity *in vitro* and a new human β-cell line has shown promising functional characteristics and can be arranged in spheroids although such approaches would lose inherent islet characteristics conferred by other cell types in the micro-organ.^[26]^

The setup developed here may also be of considerable use in exploring in detail islet function in native preparations, evaluate surrogate islets for therapy derived from iPSCs and decipher the consequences of mutations in islets of rodent or human origin.^[27]^ Indeed, most published work in this area has been done using non-physiological square-pulse stimulations with glucose jumps to supraphysiological values or, in a few cases, step-wise increases, but of glucose only. Clearly, injection of complete nutrient solutions into the animal hosting the device, or even the more ambitious administration via tube feeding, will provide a far more physiological repertoire of stimulation using the animal as a kind of bioreactor. The profiles of the electrical responses can be introduced in an FDA validated human simulator, the UVA-PADOVA T1DMS, to regulate metabolism in this in-silico model.^[15–16]^ Thus, the electrical profiling in extracorporeal islet activity, as measured here, can be transposed to regulation of glucose metabolism in humans to analyse ensuing metabolic consequences.

### 4.0 Conclusion

A large number of biosensors have been developed in the past relying on electrodes, enzymes, genetically encoded fluorescent probes or genetically modified micro-organisms.^[28]^ Cells receive various information from their environment and compute the appropriate output almost instantaneously on a millisecond scale. Although developing biosensors to harness the inherent analytic power of micro-organs or tissues faces a considerable number of technical obstacles, progress has been achieved especially in the field of bioelectronic tongues.^[29]^ To the best of our knowledge biosensors have not yet been interfaced to an entire organism with the long-term goal to achieve homeostatic control. Our work here provides a further step in the development of micro-organ based biosensors.

## 5. Experimental Section

### Animals, surgery and islet preparation

Animal experiences were conducted along ethical guidelines and authorized (APAFIS#25037-2020040917179466) by the French Ministry of Research and Innovation. Male Wistar rats (Charles River, Lyon, France), mean age 10.4 weeks and with a mean weight of 384 g were placed on a heated pad and anaesthetized with isoflurane (starting with 3.5% in air, 2 l/min; maintenance by 1.5% in air) and for analgesia meloxicam was given subcutaneously (1 mg/kg) 30 min before implantation of the catheter as well as a local anesthetic was applied (lidocaine 2,5 % cream). To insert the microdialysis catheter a small incision was placed on the interscapular area after shaving and disinfection with a povidone-iodine antiseptic (vetedine). For islet isolations, adult male C57BL/6J mice (10–20 weeks of age) were sacrificed by cervical dislocation according to University of Bordeaux ethics committee guidelines. Islets were obtained by enzymatic digestion and handpicking. MEAs were coated with Matrigel (2% v/v) (BD Biosciences, San Diego, CA) as described ^[11b, 13b, 13c]^. Islets were seeded on MEAs and cultured for 3 days at 37°C.

### Microfluidic MEA chips

For dynamic experiments, a microchannel was fabricated with PDMS-based elastomer Sylgard 184 (Dow Corning, St Denis, France) and the microchannel was fabricated by using a SU-8 mold patterned with a channel of 0.8 mm in diameter and 10 mm in length. PDMS on the mold was cured by 2 h of heating at 70°C. The microchannels where subsequently aligned on the MEAs under a microscope in a cleanroom, and bonded thanks to a previous O_2_-plasma activation. For static incubations, the chip consisted of a PDMS microwell of 3 mm in diameter and in height. Fluid shear stress was computed with the Multiphysics simulation software COMSOL 6.1 (COMSOL, Inc., Burlington, MA). First a 3D stationary simulation was carried out where the flow in the microfluidic channel is simulated following Navier Stokes equations, with a laminar flow boundary condition at the inlet. The inlet is defined in a flow rate manner, with several values: 0.2 µl/min, 1 µl/min, 5 µl/min. The shear stress is deduced from the results of the study using the following definition: T= µΔu-➔ where µ is the dynamic viscosity of water at 37°C and u-➔ is the velocity field.

### Analysis of delays in fluid transport

The delay introduced by the transit of dialysate from the subcutaneous site to its arrival in the microfluidic MEA chamber and final glucose determination after passage through the microfluidic chip and tubes was determined through analysis of videos taken with a binocular camera Moticam 5+ (Motic, Hong Kong, HK) during phenol red perfusions (stepwise gradient) under conditions identical to experimental conditions, i.e. from microdialysis to dialysate collection (see Figure 1). Phenol red solution was passed through microdialysis and the microfluidic MEA of 1 µl/min and images were taken (at 25 Hz) of the microfluidic channel and the outlet of the microfluidic MEA. Video files were analysed with an ad hoc program written in Python (Python 3.7, imageio 2.6.1, numpy 1.18.1, pandas 1.0.4) that measured a differential rate of change in colour for each region of interest (ROI). The program functions as follows: the video is first down-sampled to 10 Hz (no interpolation) and cropped to a 51×51 pixels ROI. The pixels in the ROI are spatially averaged which yields a single vector of red, green, and blue components indicating the average colour in the ROI for every frame and the red component is filtered out. The same operations are performed in a 51 x 51 pixels region of reference (ROR) where the variations in red, green and blue components are deduced from those of the ROI to produce a differential measurement that compensates for changes in ambient light. The red, green, and blue components of the differential measurement are then merged (unweighted sum) and differentiated. Finally, a moving average is applied (10 samples) for de-noising and the data are normalized (fold of maximum value). The delay from the microdialysis pump to the region of interest was measured as the peak of the calculated rate of colour change.

Data from different recording sites were synchronized to compensate for the lag introduced by the transit of dialysate through the microfluidic chip and tubes. All data were synchronized assuming t = 0 corresponds to the IP injection of glucose. The delay between blood glucose measurements and dialysate measurements due to the diffusion of glucose in the animal’s subcutaneous space (Δ*t_SQ_*) was estimated with cross-correlation analyses on experimental data.

The corrected time vector 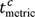 for each recorded metric is therefore, in relation to its original 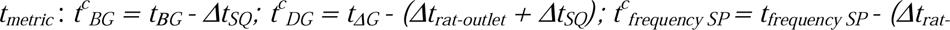 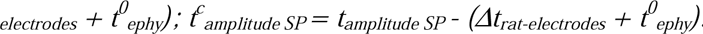 Datasets with corrected time vectors were generated with a Python script (Python 3.7, pandas 1.0.4).

### Electrophysiology

MEAs (60 MEA200/30iR-TiN, Ø 30 mm, 200 mm interelectrode distance) were purchased from Multi Channel Systems GmbH (MCS, Reutlingen, Germany). As described previously ^[12a, 13b, 13c]^, extracellular field potentials were acquired at 10 kHz, amplified, and band-pass filtered at 0.1–3,000 Hz with a USB-MEA60-Inv-System-E amplifier (MCS; gain: 1,200) or an MEA1060-Inv-BC-Standard amplifier (MCS; gain: 1,100), both controlled by MC_Rack software (v4.6.2, MCS). Electrophysiological data were analysed with MC_Rack software. SPs were isolated using a 2-Hz low-pass filter and frequencies were determined using the threshold module of MC_Rack with a dead time (minimal period between two events) of 300 ms (SPs). The peak-to-peak amplitude module of MC_Rack was used to determine SP amplitudes as published (Jaffredo et al., 2021).

### Microdialysis, glucose injections and measurements

Linear interscapular subcutaneous catheters (30 mm membrane, 20 kDa cut off; Microdialysis AB) were inserted under anaesthesia (1.5% isoflurane in air). For dialysis, Ringer dextran-60 was used (pump 107, Microdialysis AB, Kista, Sweden). For glucose or insulin tests, 2 g/kg of glucose or 2.5 U/kg of insulin were injected intraperitoneally. Blood glucose was measured in droplets collected at the caudal vein with a freestyle papillon glucometer (Abbott, Rungis, France). Glucose in the dialysate was determined using a glucose oxidase-based kit (Biolabo, Maizy, France). Human male plasma was obtained from Sigma (H4522; Sigma-Aldrich, Saint Louis, MO, USA) and contained 6 mM glucose.

### Respiration rate

The respiration rate of anaesthetised rats was measured through analysis of video files, using an ad hoc program written in Python (Python 3.7, imageio 2.6.1, numpy 1.18.1, scipy 1.6.2, pandas 1.0.4) that detects oscillating movement on the rat’s fur. The program functions as follows: the video is cropped to a ROI where movement caused by breathing is clearly visible. The video is converted to greyscale and differentiated to highlight movement. A quantity of movement is estimated for each frame by measuring the standard deviation of the differentiated pixels in the ROI. A 0.1-2.0 Hz filter is the applied to this signal and its frequency components are analysed in a spectrogram (Fast Fourier Transform, 10 s rectangular window, 0.1 s overlap). The breathing rate is retrieved by measuring the frequency of the main peak at each instant in the spectrogram through peak detection.

### Statistics

Correlations between electrophysiological data (SP frequencies and SP amplitudes) and glucose measurements (capillary and dialysate) were calculated using Spearman correlation in Python (Python 3.7, scipy 1.3.0). Electrophysiological data were resampled using windowed AUCs (see below) prior to correlation, in order to match glucose measurements. For correlation the AUC of electrophysiological data (SP frequencies and SP amplitudes) were calculated using the trapezoidal rule in time windows surrounding each glucose measurement (capillary or dialysate): AUCs used for correlation with blood glucose (BG) measurements were calculated in a [-90 s; +90 s] time window around each BG data point; AUCs used for correlation with dialysate glucose (DG) measurements were calculated in a [+0 s; +900 s] time window following each DG data point, identical to the window of dialysate collection for the corresponding DG measurement. The distinction between time windows was made because capillary blood glucose measurements were punctual (representative of a short time window), whereas the collection of dialysate samples spanned over 15 min each (representative of a 15 min time window). The diffusion delay Δt_SQ_ was estimated using cross-correlation between electrophysiological data and windowed AUCs of DG measurements. Δt_SQ_ was measured at peak correlation for each experiment individually, as it was assumed to be animal-dependent. Other statistical analyses of electrophysiological data were performed using GraphPad PRISM v7.00 (San Diego, CA, USA) with ANOVA and post-hoc tests (Dunn or Tukey) as given in the Figure legends.

## Acknowledgements

Emilie Puginier and Antoine Pirog contributed equally to this work.

We are grateful to Céline Cruciani-Guglielmacci and Christophe Magnan for helpful advices on surgery.

Fundings: This work was funded by the Agence National de Recherche [grant number ANR-18-CE17-0005 DIABLO] (to JL and SR) and a grant of the Société Francophone du Diabète (to JL).

The authors have no conflicts of interest.

