## Supplemental Material for "Continuous monitoring of glucose levels *in vivo* with a micro-organ based microfluidic biosensor"

ABSTRACT

Continuous monitoring of glucose levels has improved diabetes therapy. Current approaches rely on enzyme-linked electrochemical probes but do not allow a fully autonomous artificial pancreas. In contrast, monitoring the activity of a few electrogenic pancreatic islets in a biosensor may harness the computational power of the different endocrine cell types in the micro-organ, shaped for nutrient detection during evolution, and provide a more appropriate read-out. Extracellular electrophysiology captures slow potentials (SPs), which reflect coupled islet β-cell activity and is thus a method of choice for long-term monitoring of native islet activity *in vitro*.

We have now developed a microfluidic microelectrode chip containing a few islets and linked to interstitial fluids in live rats by subcutaneous microdialysis. The electrical activity in terms of slow potentials monitored by this biosensor reacts *ex vivo* proportionally to glucose levels off-line in serum or dialysed interstitial fluid. On-line monitoring *in vivo* reveals an excellent correlation between islet slow potential frequency, and to a lesser degree to slow potential amplitudes, to glucose concentrations with little variation between animals. The microorgan-based biosensor harness multiple parameters *in vivo* and provides a read-out closer to physiology. This demonstrates the usefulness of such biosensors for sensor-based therapy of diabetes.

**Supplementary Table 1: Spearman's ρ and p values.** Correlations and significances calculated for data presented in Figure 6 and Supplementary Figure 1.

| BLOOD GLUCOSE | | | | | | |
|---|---|---|---|---|---|---|
| Animal | Frequency vs. Amplitude | | Frequency vs. Blood Glucose | | Amplitude vs. Blood Glucose | |
| | r | 2p | r | 2p | r | 2p |
| 1 | 0,95 | 8,76E-05 | 0,93 | 0,0002 | 0,92 | 0,0005 |
| 2 | 0,95 | 2,04E-06 | 0,83 | 0,0008 | 0,84 | 0,0006 |
| 3 | 0,97 | 3,88E-07 | 0,94 | 3,93E-06 | 0,94 | 6,99E-06 |
| 4 | 0,98 | 5,41E-12 | 0,91 | 3,2E-07 | 0,96 | 1,17E-09 |
| 5 | 0,82 | 0,0021 | 0,91 | 0,0001 | 0,93 | 3,97E-05 |

| DIALYSATE | | | | | | |
|---|---|---|---|---|---|---|
| Animal | Frequency vs. Amplitude | | Frequency vs. Dialysate Glucose | | Amplitude vs. Dialysate Glucose | |
| | r | 2p | r | 2p | r | 2p |
| 1 | 0,92 | 0,0005 | 0,72 | 0,0298 | 0,78 | 0,0125 |
| 2 | 0,97 | 2E-08 | 0,95 | 5,51E-07 | 0,94 | 1,88E-06 |
| 3 | 0,99 | 1,93E-11 | 0,83 | 0,0001 | 0,88 | 1,63E-05 |
| 4 | 0,78 | 0,0125 | 0,73 | 0,0246 | 0,68 | 0,0424 |

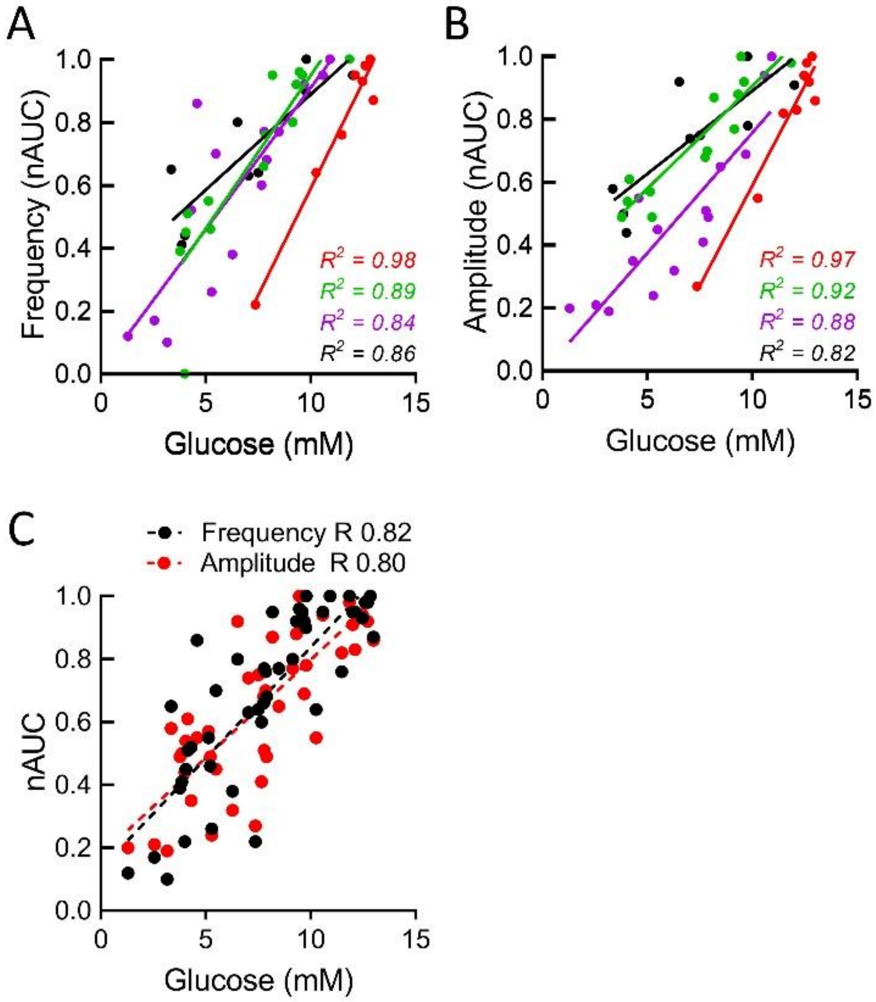


**Supplementary Figure 1: Linear correlation analysis of normalized electrical activity of islets versus dialysate glucose. A:** Correlation analysis of blood glucose levels slow potential frequencies. Each color represents one experiment (animal) and corresponding coefficients of determination $R^2$ are indicated. Data points represents normalized AUCs (1= maximal response in the corresponding experiment) for a given blood glucose value and linear regression curves are given. **B:** Correlation analysis of blood glucose levels and normalized slow potential amplitudes. Color codes as in A, corresponding $R^2$ values are indicated. **C:** Correlation analysis of slow potential frequencies (black) or amplitudes (red) at a given dialysate glucose concentration (all data points of A and B combined). Data from 4-5 animals, each 2 – 13 electrodes.

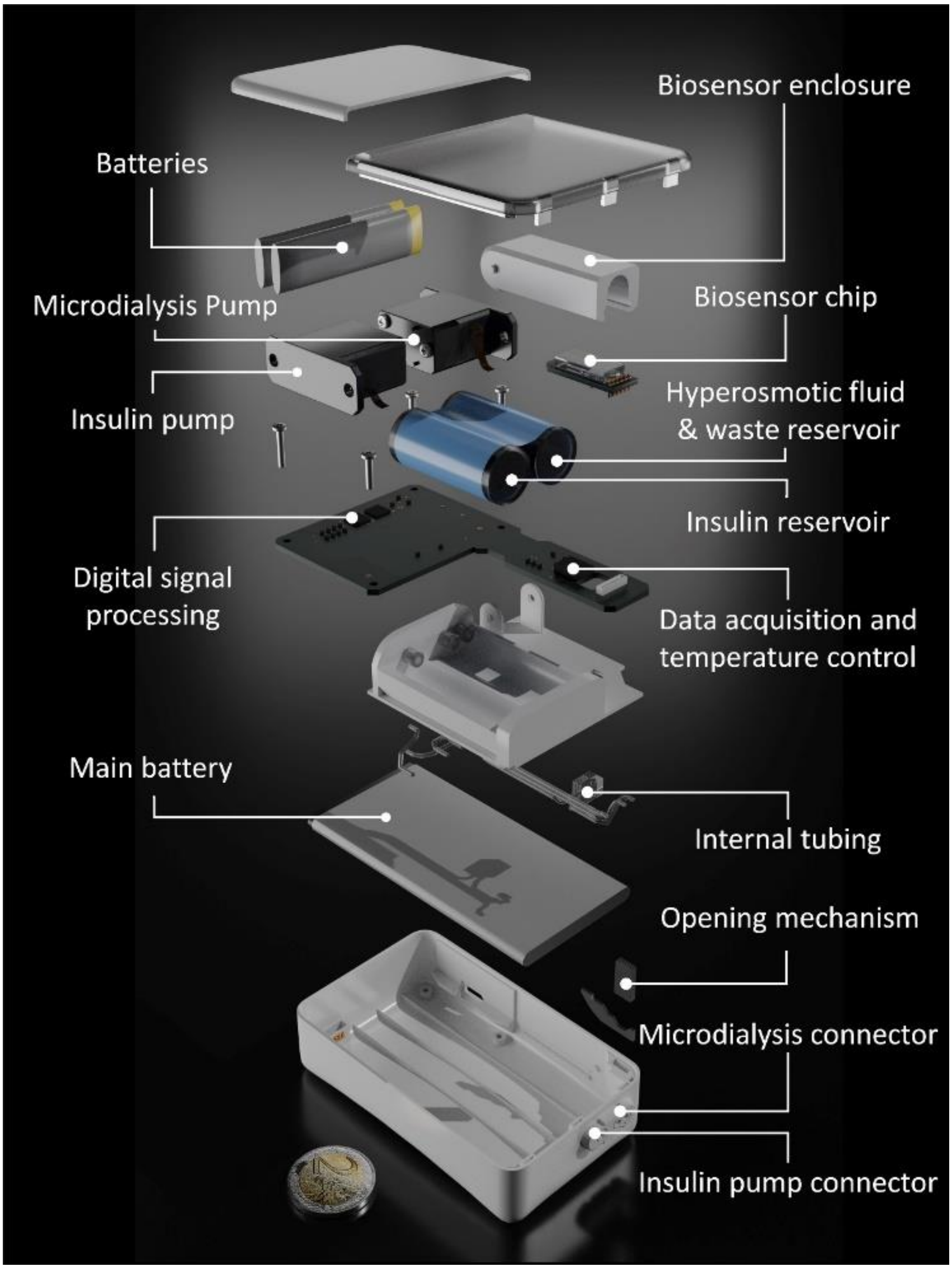


**Supplementary Figure 2: Packaging of an extracorporeal islet-based glucose monitor.** Blow-up presentation of a possible packaging of the microfluidic microelectrode device for extracorporeal use as a islet-based sensor of the demand in insulin in humans.
